# Coal-Miner: a coalescent-based method for GWA studies of quantitative traits with complex evolutionary origins

**DOI:** 10.1101/132951

**Authors:** Hussein A. Hejase, Natalie Vande Pol, Gregory M. Bonito, Patrick P. Edger, Kevin J. Liu

## Abstract

Association mapping (AM) methods are used in genome-wide association (GWA) studies to test for statistically significant associations between genotypic and phenotypic data. The genotypic and phenotypic data share common evolutionary origins – namely, the evolutionary history of sampled organisms – introducing covariance which must be distinguished from the covariance due to biological function that is of primary interest in GWA studies. A variety of methods have been introduced to perform AM while accounting for sample relatedness. However, the state of the art predominantly utilizes the simplifying assumption that sample relatedness is effectively fixed across the genome. In contrast, population genetic theory and empirical studies have shown that sample relatedness can vary greatly across different loci within a genome; this phenomena – referred to as local genealogical variation – is commonly encountered in many genomic datasets. New AM methods are needed to better account for local variation in sample relatedness within genomes.

We address this gap by introducing Coal-Miner, a new statistical AM method. The Coal-Miner algorithm takes the form of a methodological pipeline. The initial stages of Coal-Miner seek to detect candidate loci, or loci which contain putatively causal markers. Subsequent stages of Coal-Miner perform test for association using a linear mixed model with multiple effects which account for sample relatedness locally within candidate loci and globally across the entire genome.

Using synthetic and empirical datasets, we compare the statistical power and type I error control of Coal-Miner against state-of-theart AM methods. The simulation conditions reflect a variety of genomic architectures for complex traits and incorporate a range of evolutionary scenarios, each with different evolutionary processes that can generate local genealogical variation. The empirical benchmarks include a large-scale dataset that appeared in a recent high-profile publication. Across the datasets in our study, we find that Coal-Miner consistently offers comparable or typically better statistical power and type I error control compared to the state-of-art methods.

**CCS CONCEPTS:** Applied computing → *Computational genomics; Computational biology; Molecular sequence analysis; Molecular evolution; Computational genomics; Systems biology; Bioinformatics; Population genetics;*

**ACM Reference format:** Hussein A. Hejase, Natalie Vande Pol, Gregory M. Bonito, Patrick P. Edger, and Kevin J. Liu. 2017. Coal-Miner: a coalescent-based method for GWA studies of quantitative traits with complex evolutionary origins. In *Proceedings of ACM BCB, Boston, MA, 2017 (BCB),* 10 pages. DOI: 10.475/123 4

## 1 INTRODUCTION

Genome-wide association (GWA) studies aim to pinpoint loci with genetic contributions to a phenotype by uncovering significant statistical associations between genomic markers and a phenotypic trait under study. We refer to the computational methods used in a GWA analysis as association mapping (AM) methods. Among the most widely studied organisms in GWA studies are natural human populations and laboratory strains of house mouse. Recently, GWA approaches have been applied to natural populations of other organisms sampled from across the Tree of Life. For example, the study of Consortium [10] published whole genome sequences for over a thousand samples from globally distributed *Arabidopsis* populations. In combination with phenotypic data, the genomic sequence data was used in a GWA analysis to pinpoint genomic loci involved in flowering time at two different temperatures. Other recent GWA studies such as the study of Porter et al. [47] have focused on bacteria and other microbes (see [9] for a review of relevant literature).

Regardless of sampling strategy – from one or more closely related populations involving a single species to multiple populations from divergent species – it is well understood that sample relatedness can be a confounding factor in GWA analyses unless properly accounted for. Intuitively, the genotypes and phenotypes of present-day samples reflect their shared evolutionary history, or phylogeny. For this reason, covariance due to a functional relationship between genotypic markers and a phenotypic character must be distinguished from shared covariance due to common evolutionary origins. EIGENSTRAT [49] is a popular AM method which accounts for sample relatedness as a fixed effect. Other statistical AM methods have utilized linear mixed models (LMMs) to capture sample relatedness using random effects; these include EMMA [30], EMMAX [29], and GEMMA [62]. The question of whether sample relatedness is better modeled using the former or the latter – i.e., using fixed vs. random effects – is a matter of ongoing debate [50, 56].

Local variation in functional covariance across the genome is a crucial signature that AM methods use to uncover putatively causal markers. In contrast, virtually all of the most widely used state-of-the-art AM methods assume that covariance due to sample relatedness does not vary appreciably across the genome. Sample relatedness is therefore evaluated “globally” across the genome, eliding over “local” genealogical variation across loci. The latter has been observed by many comparative genomic and phylogenomic studies (see [16] for a review of relevant literature). It is well understood now that local genealogical variation within genomes is pervasive across a range of evolutionary divergence – from structured populations within a single species to multiple species at various scales up to the Tree of Life, the evolutionary history of all living organisms on Earth. Topological incongruence can be severe: for example, within a range of evolutionary conditions referred to as the “anomaly zone”, the topology of the most frequently observed local genealogy can be incongruent with the species phylogeny itself [11]. The evolutionary processes that can contribute to local genealogical variation include genetic drift and incomplete lineage sorting, recombination, gene flow, positive selection, and the combination of all of these processes (and others) [16, 18]. These observations are applicable across different GWA settings ranging from traditional studies involving closely related populations representing a single species to a comparative study involving multiple species. The latter typically involves relatively greater evolutionary divergence, which introduces added complexity in terms of accounting for sample relatedness.

Computational approaches for detecting local genealogical variation are broadly characterized by their modeling assumptions. One class of methods makes use of the Four-Gamete Test [24], which requires the simplifying assumption that sequence evolution can be described by the infinite sites model. Methods in this class include the LRScan algorithm [59]. Another class of parametric methods make use of finite sites models of sequence evolution. One example is RecHMM [61], which applies a sequentially Markovian approximation [40] to the full coalescent-with-recombination model [21]. More recently, coalescent-based methods have been developed to infer local coalescent histories and explicitly ascribe local genealogical variation to different evolutionary processes. Examples include Coal-HMM [15, 20, 37, 38] and PhyloNet-HMM [35].

To address this methodological gap, we recently introduced Coal-Map, a new AM method that accounts for local genealogical variation across genomic sequences. Coal-Map performs statistical inference under a linear mixed model (LMM). The LMM utilizes fixed effects to account for global sample relatedness and, depending upon whether the test marker is located within a locus containing putatively causal markers, local sample relatedness as well. The latter condition is evaluated using model selection criteria. Coal-Map required local-phylogeny-switching breakpoints as input. We validated Coal-Map’s performance using simulated and empirical data. Our performance study demonstrated that Coal-Map’s statistical power and type I error control was comparable or better than other state-of-the-art methods that account for sample relatedness using fixed effects.

## 2 METHODS

### 2.1 Overview of Coal-Miner algorithm

We begin by introducing the high-level design of Coal-Miner, our new algorithm for statistical AM which accounts for local variation of sample relatedness across genomic sequences. The input to the Coal-Miner algorithm consists of: (1) an *n* by *k* multi-locus sequence data matrix *X*, (2) a phenotypic character *y*, and (3) *ℓ*^*^, the number of “candidate loci” used during analysis, where a candidate locus is a locus that is inferred to contain one or more putatively causal SNPs. The output consists of an association score for each site *x* ∈ *X*.

Coal-Miner’s statistical model captures the relationship between genotypic data *X* and the phenotypic character *y* in the form of a linear mixed model (LMM). The LMM incorporates multiple effects to capture the phenotypic contributions of and local genealogical variation among multiple candidate loci. A candidate locus is represented by a fixed effect, and a random effect is included to capture “global” sample relatedness as measured across all loci in *X*. Ideally, the set of candidate loci identified during a Coal-Miner analysis is identical to the set of causal loci (i.e., loci containing causal SNPs) for the trait under study; in practice, the set of candidate loci are inferred as part of the Coal-Miner algorithm, which we discuss in greater detail below. The LMM takes the following form (in the notation of Zhou and Stephens [62]):

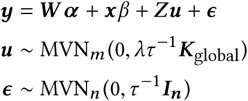

The fixed effects are represented by covariates *W* with coefficients *α*, which include covariates that capture local sample relatedness within each candidate locus, and the test SNP *x* with effect size *β*. Global sample relatedness (i.e., sample relatedness as measured across all loci in the genotypic data *X*) is specified by the relatedness matrix *K* global. The random effects *u* and *ϵ* account for global sample relatedness and residual error, respectively. Each of the two random effects follows an *m* - *dimensional* multivariate normal distribution (abbreviated “MVN”) with mean 0. The random effects *u* have covariance *λτ* ^-1^*K* global and the random effects *ϵ* have covariance *τ* ^-1^*I*_*n*_, where *λ* is the relative ratio between the two, *I*_*n*_ is the identity matrix, and the residual errors have variance *τ* ^-1^. *Z* is the design matrix corresponding to random effects *u*.

The design of the Coal-Miner algorithm takes the form of a methodological pipeline. We now discuss each pipeline stage in turn.

### 2.2 Stage one of Coal-Miner: inferring local-phylogeny-switching breakpoints

The input to the first stage of Coal-Miner is the genotypic data matrix *X*. The output consists of a set of local-phylogeny-switching breakpoints *b* which partition the sites in *X* into loci {*X*_*i*_}, where 1 ≤ *i* ≤ ℓ and ℓ is the number of loci. We require that ℓ≤ ℓ. (The ratio of ℓ^*^ and ℓ depends upon the genomic architecture of the trait corresponding to character *y*.)

The general approach to address this computational problem is to infer local coalescent histories under an appropriate extension of the multi-species coalescent (MSC) model [22, 31, 58], and then to assign breakpoints based upon gene tree discordance. Each pair of neighboring breakpoints delineates a locus for use in downstream stages of the Coal-Miner pipeline. The specific choice of model/method depends upon the relevant evolutionary processes involved in multi-locus sequence evolution, particularly regarding the source(s) of local genealogical discordance.

In this study, we use one of two different methods, depending upon assumptions about biomolecular sequence evolution. In the simulation study, the simulations make use of the infinite sites model. We therefore used the LRScan algorithm [59] to compute local-topology-switching breakpoints based upon the Four Gamete Test (FGT) [24]. In the empirical study, we did not make use of the infinite sites model and its assumptions about sequence evolution. Furthermore, multiple evolutionary processes were known to be involved in multi-locus sequence evolution, including genetic drift/incomplete lineage sorting (ILS), recombination/gene conversion, gene flow/horizontal gene transfer (HGT), and natural selection. Breakpoint inference under the corresponding extended MSC model is suspected to be a computationally difficult problem. Existing methods for this problem (e.g., PhyloNet-HMM [35]) did not have sufficient scalability for the dataset sizes examined in our study. As a more feasible alternative, we inferred local-topology-switching breakpoints using Rec-HMM [61]. Rec-HMM performs fixed-species-phylogeny inference of local genealogies under a statistical model that combines a finite-sites substitution model and a hidden Markov model which is meant to capture intra-sequence dependence (such as arises from recombination and other evolutionary processes).

### 2.3 Stage two of Coal-Miner: identifying candidate loci

The input to the second stage of Coal-Miner consists of the genotypic data matrix *X*, the set of breakpoints *b* which partition *X* into loci {*X*_*i*_}, where 1 ≤ *i* ≤ ℓ and ℓ is the number of loci, the phenotypic character *y*, and ℓ^*^, the number of candidate loci to identify. The output is a set of candidate loci 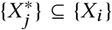 where 1 ≤*j* ≤ ℓ^*^.

Our general approach to this problem consists of a search among possible sets of candidate loci 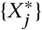 using optimization under a “null” version of Coal-Miner’s LMM, where we do not consider a test SNP (i.e., *β* = 0 in Coal-Miner’s LMM) and the phenotypic contributions from causal SNPs in each candidate locus 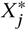 is captured by covariates {*w*_*j*_} ⊆ *W*. Since we compare fitted LMMs that may have varying fixed effects, we use LMM log-likelihood as our optimization criterion (reproduced from equation (3) in [62]):

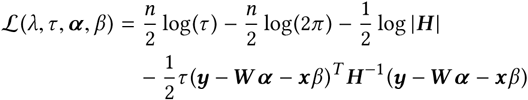

where *G* = *Z K*_global_*Z*^*T*^ and *H* = *λG* +*I*_*n*_. Due to the computational difficulty of this optimization problem, numerical optimization procedures are typically used. We obtained estimates of *λ* in the range of [10^−5^, 1] using the optimization heuristic implemented in the GEMMA software library [62], which combines Brent’s method [8] and the Newton-Raphson method.

For each candidate locus 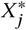, sample relatedness was evaluated using principal component analysis (PCA) [27] of 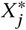 – similar to techniques that are widely used by AM methods to account for global sample relatedness as fixed effects [49]. The phenotypic contribution of candidate locus 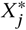 was represented using covariates {*w*_*j*_} which consisted of the top five principal components. (The *z*th principal component corresponds to the sample covariance matrix eigenvector with the *z*th largest eigenvalue.) For added computational efficiency, we substituted the following search heuristic in place of set-based search among all possible ℓ^*^-size sets of candidate loci. For each locus *X*_*i*_, we used MLE to fit an equivalent LMM, except that the covariates *W* included only the covariates {*w*_*i*_} for locus *X*_*i*_(as computed using the above PCA-based procedure). The output set of candidate loci consists of the top ℓ^*^ loci based upon fitted LMM likelihood.

### 2.4 Stage three of Coal-Miner: SNP-based association testing

The input to the third stage of Coal-Miner consists of the genotypic data matrix *X*, the set of breakpoints *b* which partition *X* into loci {*X*_*i*_}, where 1 ≤ *i* ≤ ℓ and ℓ is the number of loci, the phenotypic character *y*, and the set of candidate loci 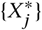. The output of this stage is Coal-Miner’s final output.

Each test SNP *x* is tested for association under Coal-Miner’s LMM. Variation in local sample relatedness across candidate loci 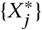 is captured by covariates in *W*: specifically, if the test SNP *x* is located within a candidate locus 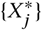, the covariates *W* include a corresponding covariate *w*_*j*_which consists of the top principal component from PCA applied to 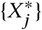 (see above discussion of previous stage), and otherwise not. (Stages two and three of the Coal-Miner pipeline utilize different covariates *W* due to the absence or presence of a test SNP effect in their respective LMMs.) The LMM is fitted using the likelihood-based numerical optimization procedures that are also used in stage two of Coal-Miner, and the association score is computed using a likelihood ratio test.

### 2.5 Simulation study

#### Experiments involving quantitative traits with varying genomic architectures

Neutral simulations of multi-locus sequence data were based upon either tree-like or non-tree-like evolutionary scenarios. The evolutionary scenarios shared a species phylogeny that we used in a prior simulation study (shown in Supplementary Figure S1 panels (a) and (b)). We used ms [23] to simulate coalescent histories (and embedded gene trees) under an extension of the coalescent model [31] which allows instantaneous unidirectional admixture (IUA) [14]. Under this model, the parameterization of the model phylogeny includes an admixture proportion *γ*. Appropriate choices of *γ* allow us to explore the impact of tree-like and non-tree-like evolution in our simulation study, where we utilized a *γ* of 0.0 and 0.5, respectively. Each replicate dataset sampled 10 independently and identically distributed loci and 1000 individuals; taxa A, B, and C were represented by 250, 250, and 500 samples, respectively. Bi-allelic sequence evolution was simulated under the infinite sites model to obtain 250 bp per locus, resulting in total sequence length of 2.5 kb per replicate dataset.

As a means to investigate the impact of the genomic architecture of phenotypes, we simulated phenotypic characters using the approach from our previous work [19]. For each synthetic multi-locus sequence dataset in the neutral simulations, we randomly selected either 10%, 20%, or 30$ of loci as causal. Twenty causal SNPs were then randomly selected from causal loci such that each causal locus contained at least one causal SNP and causal SNPs had minor allele frequency between 0.1 and 0.3. Given a set of causal SNPs *δ*, we sampled character *y* under an extension of the quantitative trait model used by Long and Langley [36] and Besenbacher et al. [6]. The trait value for the *i*th individual is represented as:

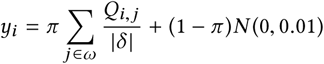

where *π* specifies the ratio between the genotypic contribution and an environmental residual, *Q* is 1 if sample *i* has the derived allele at the *j*th causal SNP and 0 otherwise, and the environmental residual is normally distributed with mean 0 and standard deviation 0.01. Our simulations utilized a ratio *π* of 0.5.

Our simulation study also included non-neutral simulations that incorporated positive selection. We used msms [17] to conduct forward-time coalescent simulations of genotypic sequence evolution (in place of an otherwise equivalent neutral backward-time coalescent simulation using ms), where causal loci were evolved under deme-dependent positive selection with a finite sites mutation model and all other loci evolved neutrally (as discussed above in the neutral simulation procedure). We used a selection coefficient of *s* = 0.56, which is in line with estimates from prior studies of positive selection in natural *Mus* populations [54]. Qyantitative traits with between one and three causal loci were simulated using the above procedure.

The simulation study experiments involving quantitative traits with varying genomic architectures included 12 different model conditions in total. To recap, the model conditions differed in terms of the number of causal loci (between one and three), model phylogeny (either tree-like or non-tree-like), and the presence or absence of positive selection. For each model condition, we repeated the simulation procedure to obtain 20 replicate datasets.

#### Experiments involving alternative evolutionary scenarios

Multi-locus sequence evolution in the above simulations is impacted by genetic drift and incomplete lineage sorting, admixture, positive selection, and combinations of these processes. Our simulation study also included additional model conditions that involved alternative models of multi-locus sequence evolution. Each model condition was an extension of the above neutral model condition with 10% causal loci. One set of model conditions varied split time *t*_1_ in the above model tree (i.e., *γ* = 0). Another set of model conditions varied admixture time *t*_1_ in the above model phylogeny where *γ* = 0.5. The impact of recombination was explored in a model condition which made use of the coalescent-with-recombination model [21]. The simulations generated 2.5 kb alignments under a finite-sites model of recombination with per-generation crossover probability between adjacent sites of 10^−9.85^, which is 1-2 orders of magnitude smaller than estimates for mouse, rat and human [26]. We further explored the impact of gene flow using a model condition which substituted the isolation-with-migration model [44] in place of the IUA model.

#### Open data access, methods, and performance evaluation

Detailed software commands and instructions for accessing simulation study datasets under an open license are provided in the SI.

The other methods in our study consisted of Coal-Map, GEMMA, and EIGENSTRAT. We followed the procedure from our original study [19] to obtain FGT-based local-phylogeny-switching breakpoints and run Coal-Map analyses. For consistency with the other LMM-based AM methods in our study, we ran GEMMA using an IBS kinship matrix as our measure of global sample relatedness and MLE and LRT to obtain association scores. EIGENSTRAT was run with default settings using the top ten principal components from the genotypic data matrix *X*, following the recommendations of Price et al. [49]. Detailed software commands are listed in the SI.

We evaluated performance based on statistical power, type I error, and AUROC. To compare AUROC, we performed Delong *et al.* tests [12] using the Daim v. 1.1.0 package [48] in R [51]. Custom scripts were used to conduct the simulation study; all scripts are provided under an open source license (see SI for details and download instructions).

### 2.6 Empirical study

#### Arabidopsis dataset

The dataset consists of whole genome sequence (WGS) data and phenotypic data for two quantitative traits: flowering time at 10 °C and 16 °C. A total of 1,135 samples from natural populations across the globe are represented. The phylogeny shown in Supplementary Figure S16 depicts the geographic origins of and evolutionary relationships among the samples. The dataset was originally published and analyzed by Consortium [10], and we obtained genomic sequences and quantitative trait data from the 1001 Genomes Project database [10] (accessible at www.1001genomes.org); the former includes both assembled WGS data and variant calls for a total of 10,707,430 biallelic SNPs. (Details about sequencing, assembly, filtering, quality controls, and variant calling are described in [10].)

Stage one of the Coal-Miner pipeline made use of RecHMM [61] to infer local-phylogeny-switching breakpoints. For computational efficiency, the breakpoint inference utilized a subset of taxa rather than the full set of taxa. The subset was chosen to maximize evolutionary divergence and was comprised of one sample from each of the following geographic regions: Spain, Sweden, USA, and Russia. For chromosomes 1 through 5, the analysis in stage one resulted in 1876, 991, 783, 559, and 913 loci with an average locus length of 16 kb, 19 kb, 30 kb, 33 kb, and 29 kb, respectively.

Using the loci obtained in stage one as input, the second stage of Coal-Miner was run on both trait characters. The 10 °C analysis identified 179, 99, 108, 109, and 95 candidate loci in chromosomes 1 through 5, respectively. The 16 °C analysis identified 115, 42, 88, 65, and 89 candidate loci in chromosomes 1 through 5, respectively. Coal-Miner also requires that ℓ^*^, the number of candidate loci, be provided as an input parameter. We followed the general approach of Solís-Lemus and Ané[53] to determine a suitable value for ℓ^*^. Specifically, we calculated the likelihood score of the fitted “null” LMM for each locus (see above), and we examined the distribution of likelihood scores (Supplementary Figure S15). We then assigned ℓ^*^ based on the distribution’s inflection point.

The inputs to the third stage of Coal-Miner consisted of the set of candidate loci, a quantitative trait character (flowering time at either 10 °C or 16 °C), and the genotypic sequence data matrix which consisted of sites with minor allele frequency threshold of 0.3. The third stage of Coal-Miner was run using the same settings as in the simulation study.

#### Heliconius erato dataset

We re-analyzed data from the study of Supple et al. [57]. The dataset includes 45 *H. erato* samples collected from four hybrid zones located in Peru, Ecuador, French Guiana, and Panama. Each sample exhibits one of two red phenotypes – postman and rayed – where 28 samples had the postman phenotype and 17 samples had the rayed phenotype. The genotypic data were sequenced from the 400 kb genomic region referred to as the D interval in *H. erato*. The D interval spans 56,862 biallelic SNPs and is known to modulate red phenotypic variation. Coal-Miner was run on the *H. erato* dataset using the same approach as in the *Arabidopsis* dataset analysis (see above). The first stage of Coal-Miner identified seven loci and the second stage inferred a single candidate locus.

#### Burkholderiaceae dataset

Bacteria belonging to the *Burkholderiaceae* are of interest given their importance in human and plant disease, but also given their role as plant and fungal endosymbionts and their metabolic capacity to degrade xenobiotics. Fully sequenced (closed) genomes belonging to *Burkholderiaceae* were selected and downloaded from the PATRIC web portal (www.patricbrc.org/) [60]. Supplementary Table S5 lists sampled species names along with other information (IDs, groups, and pathogenicity). We chose to maximize phylogenetic and ecological diversity in this sampling, so we included available genomes belonging to free-living, pathogenic, and endosymbiotic species spanning across the genera *Burkholderia*, *Ralstonia*, *Pandoraea*, *Cupriavidus*, *Mycoavidus*, and *Polynucleobacter*. A total of 57 samples were included, of which 52 samples were free-living and 5 were endosymbionts. Genomes ranged in size from 1.56 Mb to 9.70 Mb and spanned between 2,048 and 9,172 coding DNA sequences (CDS). The software package Proteinortho [32] was run using default parameters to detect single copy orthologs in the selected genomes. A total of 549 orthologs were recovered in the Proteinortho analysis. We analyzed a phenotype that identified each sample’s status as either an animal pathogen or non-animal pathogen. Coal-Miner was used to analyze the genomic sequence data and phenotypic character using the same approach as in the other empirical analyses (see above). The initial stages of Coal-Miner identified 55 candidate loci. Genes with significant associations based upon the Coal-Miner analysis were further classified based upon their Gene Ontology [3] and KEGG [28] pathway assignments.

## 3 RESULTS

In this study, we introduce Coal-Miner, a new statistical AM method which advances the state of the art in terms of its statistical power and type I error control. Coal-Miner’s performance advantage derives primarily from two factors. First, Coal-Miner utilizes a new LMM with multiple effects to explicitly capture the genomic architecture of a phenotype, where both genotypic and phenotypic characters are the product of a complex evolutionary history which can cause sample relatedness to vary locally across genomic loci. The LMM captures global sample relatedness as a random effect, in contrast to the fixed-effect approach used by Coal-Map. Second, the pipeline-based design of Coal-Miner incorporates an intermediate stage to infer candidate loci for use in the new LMM. We validated the performance of Coal-Miner using two different types of datasets: synthetic datasets and empirical datasets sampled from natural populations of non-model organisms – each from a different kingdom. The synthetic datasets were simulated under a range of evolutionary scenarios that included multiple causes of local genealogical variation (see Methods for more details).

## 3.1 Simulation study

### Experiments involving varying genomic architecture of a quantitative trait

We conducted experiments that varied the proportion of causal loci as a means to investigate the impact of the genomic architecture of a trait on AM method performance. The model conditions utilized simulations with between 10% and 30% causal loci and either neutral or non-neutral evolution on either tree-like or nontree-like model phylogenies. The methods under study included Coal-Miner, our new AM method, as well as representative methods from different classes of state-of-the-art methods: Coal-Map, a LMM-based AM method that accounts for local and global sample relatedness as fixed effects, GEMMA, a LMM-based AM method that accounts for global sample relatedness as a random effect (but does not account for local sample relatedness), and EIGENSTRAT, an AM method that accounts for global sample relatedness as a fixed effect (but does not account for local sample relatedness). We compared the statistical power and type I error control of each method using receiver operating characteristic (ROC) curves (Supplementary figures S2 through S5), and Table 1 1 compares the area under ROC curve (AUROC) of each method.

**Table 1:** The impact of the genomic architecture of a quantitative trait on the performance of Coal-Miner and the other AM methods. Multi-locus sequences were simulated under neutral or non-neutral evolution on tree-like or non-tree-like model phylogenies, and quantitative traits were simulated using causal markers sampled from 10%, 20%, or 30% of loci (see Methods section for more details). The performance of each AM method was evaluated based on the area under its receiver operating characteristic (ROC) curve, or AUROC. We report each method’s average AUROC across twenty replicate datasets for each model condition. Coal-Miner’s AUROC is shown in bold where it significantly improved upon the AUROC of the most accurate of the other AM methods, based upon the test of DeLong et al. [12] (*n* = 20; *α* = 0.05). We corrected for multiple tests using the approach of Benjamini and Hochberg [5], and corrected q-values are shown. (The corresponding ROC plots are shown in Supplementary Figures S2 through S5.)

| Model condition |  |  | AUROC |  |  |  | q-value |
| --- | --- | --- | --- | --- | --- | --- | --- |
| Neutral vs. non-neutral | Model phylogeny | Percentage of causal loci (%) | Coal-Miner | Coal-Map | GEMMA | EIGENSTRAT |  |
| Neutral | Non-tree-like | 10 | **0.962** | 0.939 | 0.866 | 0.871 | < 0.00001 |
|  |  | 20 | **0.921** | 0.899 | 0.849 | 0.859 | < 0.00001 |
|  |  | 30 | **0.904** | 0.882 | 0.847 | 0.832 | < 0.00001 |
| Neutral | Tree-like | 10 | **0.943** | 0.922 | 0.87 | 0.833 | 0.0053 |
|  |  | 20 | **0.904** | 0.847 | 0.843 | 0.813 | < 0.00001 |
|  |  | 30 | **0.904** | 0.853 | 0.844 | 0.799 | 0.00003 |
| Non-neutral | Non-tree-like | 10 | **0.959** | 0.933 | 0.896 | 0.836 | < 0.00001 |
|  |  | 20 | **0.926** | 0.897 | 0.856 | 0.847 | < 0.00001 |
|  |  | 30 | **0.894** | 0.863 | 0.832 | 0.816 | < 0.00001 |
| Non-neutral | Tree-like | 10 | **0.954** | 0.922 | 0.856 | 0.841 | < 0.00001 |
|  |  | 20 | **0.89** | 0.85 | 0.832 | 0.796 | 0.00003 |
|  |  | 30 | **0.879** | 0.836 | 0.83 | 0.783 | 0.0007 |

Regardless of the proportion of causal loci and the evolutionary scenario explored in these model conditions, Coal-Miner’s AUROC was significantly better than the next best method in our study (either Coal-Map or GEMMA) based upon the corrected test of DeLong et al. [12] (Table 1). A similar observation was made when measuring performance using true positive rate (TPR) at a false positive rate (FPR) of 0.1 (Supplementary Table S2), except that Coal-Miner’s performance advantage over the next best method was even more pronounced. The TPR difference was 0.158 on average and ranged as high as 0.248. Across these model conditions, we observed a consistent ranking of AM methods by AUROC (with two minor exceptions): Coal-Miner first, Coal-Map second, GEMMA third, and EIGENSTRAT fourth. The minor exceptions involved the two lowest AUROC values on the neutral, non-tree-like model condition with 10% or 20% causal loci, where GEMMA and EIGENSTRAT swapped rankings. We noted that Coal-Map’s AUROC was second best on model conditions with the smallest proportion of causal loci, but its performance tended to degrade as the proportion increased. Coal-Map’s AUROC was only marginally better than GEMMA on model conditions with the highest proportion of causal loci.

The impact of varying the proportion of causal loci was similar for all methods: AUROC tended to degrade as the proportion of causal loci increased from 10% to 30%. However, Coal-Miner’s performance advantage relative to the other AM methods was flat or improved as the proportion of causal loci increased.

The model conditions included different combinations of genetic drift/incomplete lineage sorting and/or gene flow – evolutionary processes which can generate local variation in sample relatedness. Note that model conditions with non-tree-like model phylogenies incorporated all of these evolutionary processes (including genetic drift/incomplete lineage sorting). The impact of the different evolutionary processes differed across the methods. Coal-Miner’s AUROC tended to be larger on model conditions involving both drift/ILS and gene flow as sources of local genealogical variation, and Coal-Map’s AUROC was similarly affected. On the other hand, GEMMA’s AUROC was comparable (within 0.01) based on this comparison, with the exception of non-neutral model conditions involving 10% or 20% causal loci.

A comparison of model conditions that differed only with respect to neutral versus non-neutral evolution revealed the impact of positive selection on AM method performance. We note that, in our experiments, causal loci evolved differentially compared to non-causal loci since positive selection acted only upon the former but not the latter. Coal-Miner and Coal-Map returned comparable AUROC (within 0.025) regardless of neutral versus non-neutral evolution. GEMMA and EIGENSTRAT performed similarly, although slightly greater variability (within 0.035) was observed. For LMM-based methods, there was no obvious trend in terms of direction of change when comparing neutral versus non-neutral experiment results. There was an apparent trend for EIGENSTRAT, however: positive selection tended to reduce EIGENSTRAT’s AUROC, with one exception (model conditions with a tree-like model phylogeny and 10% causal loci).

### Experiments involving alternative evolutionary scenarios

Our simulation study also included additional experiments that explored other neutral evolutionary scenarios. These model conditions fixed the proportion of causal loci to 10%. Supplementary Table S1 shows an AUROC comparison of Coal-Miner and the other AM methods on the additional model conditions.

For model conditions that varied divergence time, involved recombination, or incorporated an isolation-with-migration (IM) model of gene flow, Coal-Miner returned significantly improved AUROC compared to the next best method based upon the test of DeLong et al. [12], and the other AM methods were ranked similarly to the experiments which varied the proportion of causal loci. A similar ranking was obtained when performance was measured using TPR at an FPR of 0.1 (Supplementary Table S3). Coal-Miner returned a comparable AUROC (within 0.027) as the divergence time *t*_1_ increased from 1.0 to 2.9. The other methods performed similarly, except that the AUROC difference was larger (within 0.031). In the IM-based model condition, all methods returned AUROC that was comparable relative to experiments using the IUA model that were otherwise equivalent.

For IUA-based model conditions that varied the admixture time *t*_1_, Coal-Map and Coal-Miner had comparable AUROC which was better than GEMMA and EIGENSTRAT. When comparing TPR at an FPR of 0.1, Coal-Miner returned a significant performance improvement relative to Coal-Map and the other AM methods (Supplementary Table S3). As seen in Supplementary Figures S8 and S9, Coal-Miner’s TPR was better than Coal-Map when the false positive rate was 0.1 or less; the reverse was true only for large false positive rates (greater than around 0.15 for the *t*_1_ = 1.0 model condition and greater than around 0.2 for the *t*_1_ = 2.9 model condition). Among the AM methods in our study, Coal-Miner’s AUROC was least impacted by the choice of admixture time and differed by at most 0.029 as the time *t*_1_ increased from 1.0 to 2.9. The AUROC of the other AM methods became smaller as the admixture time became more ancient, and the AUROC difference was relatively greater than Coal-Miner (as much as 0.086).

## 3.2 Empirical study

To demonstrate the flexibility of the Coal-Miner framework, we conducted Coal-Miner analyses of three empirical datasets which spanned a range of GWAS settings. Each of the three datasets sampled taxa from a different kingdom and ranged from well-studied organisms to relatively novel organisms about which little is known. Specifically, the datasets sampled (1) natural populations of a single plant species, (2) multiple closely related butterfly species where gene flow is a countervailing force versus genetic isolation, and (3) divergent bacterial species where horizontal gene transfer is suspected to be rampant. The datasets also varied in terms of the evolutionary processes with first-order impacts upon genome/phenotype evolution. The empirical analyses served two purposes: methodological validation using positive and negative controls based upon previous literature, and generation of new hypotheses for future study.

### Arabidopsis dataset

We used Coal-Miner to re-analyze an *Arabidopsis* dataset which the 1001 Genomes Consortium published in Cell this past summer [10]. The dataset includes samples from 1,135 high quality re-sequenced natural lines adapted to different environments with varying local climates [2, 10]. The sampled data included whole genome sequences and quantitative trait data for two traits: flowering time under high and low temperature – 16 °C and 10 °C, respectively.

A key component of the study of the 1001 Genomes Consortium was a GWA analysis of the genomic sequences and quantitative trait data using EMMAX [29], another state-of-the-art statistical AM method (see [62] for a comparison of EMMAX and other stateof-the-art statistical AM methods examined in our study). A major focus of the analysis was a set of five genes which are known to regulate flowering and contribute to flowering time variation at 10 °C in *Arabidopsis* [2, 25, 41]: FLOWERING LOCUS T (FT), SHORT VEGETATIVE PHASE (SVP), FLOWERING LOCUS C (FLC), DELAY OF GERMINATION 1 (DOG1), and VERNALIZATION INSENSITIVE 3 (VIN3). Plants rely on both endogenous and environmental (e.g. temperature and photoperiod) cues to initiate flowering [1, 2]. These five genes encode major components of the vernalization (exposure to the prolonged cold) and autonomous pathways known to regulate the initiation of flowering in *Arabidopsis*. Allelic and copy number variants (CNV) for many of these genes, including FLC, are known to serve important roles in generating novel variation in flowering time and permit plants to adapt to new climates [39, 42, 45].

Under a conservative Bonferroni-corrected threshold [7], CoalMiner identified significant peaks associated with flowering time under high and low temperature (16 °C and 10 °C, respectively). In particular, Coal-Miner identified significantly associated markers in all five genes (FT, SVP, FLC, DOG1, and VIN3) for both the 16 °C dataset and the 10 °C dataset (Supplementary Figure S12). Within the five genes, Coal-Miner analyses returned peaks which largely agreed across the 10 °C and 16 °C datasets. Some differences involved association scores that were borderline significant in one dataset but not the other.

Table 2 compares the Coal-Miner analysis with similar analyses using two other state-of-the-art statistical AM methods. The EMMAX analysis in the study of Consortium [10] identified significant associations for three of the genes at 10 °C, and association score peaks were marginally below a Bonferroni-corrected threshold in the other two genes (SVP and FLC). Furthermore, significant peaks were only detected in DOG1 at 16 °C, but no significant peaks were detected in the other four genes for this dataset. DOG1 is known to be involved in determining seasonal timing of seed germination and influences flowering time in Arabidopsis [25]. (See Figure 2 in [10] for the original Manhattan plot.) GEMMA’s performance was qualitatively similar to EMMAX (Supplementary Figure S13). At 10 °C, GEMMA recovered significant associations in three of the genes but not in the remaining two (SVP and FLC); at 16 °C, no significant peaks were detected in three genes, a peak just above the threshold of significance was detected in FT, and another peak was detected in DOG1.

**Table 2:** A comparison of Coal-Miner and two other state-of-the-art statistical AM methods based upon analyses of the two *Arabidopsis* datasets. The other AM methods are GEMMA and EMMAX, the statistical AM method used in the study of Consortium [10]. We evaluate whether the three AM methods detected significantly associated markers in five genomic regions centered on positive control genes which are known to regulate flowering time in *Arabidopsis*. We use a Bonferroni-corrected threshold for significance. For two of the five genomic regions in the 10 °C dataset, EMMAX returned association scores that were near the threshold of significance (marked using an asterisk). The corresponding Manhattan plots for the Coal-Miner and GEMMA analyses are shown in Supplementary Figures S12 and S13, respectively. The corresponding Manhattan plot for the EMMAX analysis is shown as Figure 2 in [10].

| Dataset | Positive control gene | Significantly associated markers detected? |  |  |
| --- | --- | --- | --- | --- |
|  |  | Coal-Miner | EMMAX | GEMMA |
| 10 °C | FLOWERING LOCUS T (FT) | Yes | Yes | Yes |
| 10 °C | SHORT VEGETATIVE PHASE (SVP) | Yes | No* | No |
| 10 °C | FLOWERING LOCUS C (FLC) | Yes | No* | No |
| 10 °C | DELAY OF GERMINATION 1 (DOG1) | Yes | Yes | Yes |
| 10 °C | VERNALIZATION INSENSITIVE 3 (VIN3) | Yes | Yes | Yes |
| 16 °C | FLOWERING LOCUS T (FT) | Yes | No | Yes |
| 16 °C | SHORT VEGETATIVE PHASE (SVP) | Yes | No | No |
| 16 °C | FLOWERING LOCUS C (FLC) | Yes | No | No |
| 16 °C | DELAY OF GERMINATION 1 (DOG1) | Yes | Yes | Yes |
| 16 °C | VERNALIZATION INSENSITIVE 3 (VIN3) | Yes | No | No |

### Heliconious dataset

Supplementary Figure S14 displays the Manhattan plot generated after applying Coal-Miner on the *H. erato* dataset across the D interval. We identified two significant peaks ranging from 502 kb to 592 kb and 658 kb to 682 kb, respectively. The second peak is located at the 3, of the optix transcription factor, a gene previously shown to be behind the red phenotype variation in *Heliconius* [57]. The first peak is located in a noncoding region more distant from the 3, of the optix transcription factor.

### Burkholdericeae dataset

We applied Coal-Miner on an empirical dataset of complete genomes of bacteria belonging to the *Burkholderiaceae* and spanning a diversity of ecological states including animal and plant pathogens. Supplementary Table S4 shows the genes inferred by Coal-Miner to be associated with human pathogenicity, along with their inferred KEGG pathway and gene ontology assignments. In total, we identified 16 genes associated with human pathogenicity in *Burkholderia*. Four of these genes have been implicated in pathogenicity by others, and in some cases validated through gene knockout and experimental evolution experiments. For example, the cell division protein FtsK that Coal-Miner associated with human pathogenicity was found to be one of three genes under positive selection in *Burkholderia multivorans* during a 20-year cystic fibrosis infection [52]. Modifications of another gene identified by Coal-Miner, DNA gyrase subunit A, are well known to be implicated with virulence and antibiotic resistance to quinolone and ciprofloxacin in pathogenic *Burkholderia* [4, 55]. For example, Lieberman et al. [34] found that the DNA gyrase subunit A gene was under positive selection during a *Burkholderia dolosa* outbreak among multiple patients with cystic fibrosis [34]. Another gene identified by Coal-Miner, Excinuclease ABC subunit A, has been shown to bind to previously published vaccine targets [43]. Coal-Miner also associated the protein dihydrofolate synthase with animal pathogenicity. Point mutations leading to nonsynonymous base changes in the dihydrofolate reductase gene have previously been demonstrated to be associated with trimethoprim resistance in cystic fibrosis patients infected by *Burkholderia cenocepacia* [13, 33].

## 4 DISCUSSION

### 4.1 Simulation study

For the model conditions that varied the proportion of causal loci with neutral or non-neutral evolution on tree-like or non-tree-like model phylogenies, Coal-Miner had better performance than all of the other state-of-the-art methods in our study, as measured using AUROC and TPR at an FPR of 0.1. This suggests that Coal-Miner’s performance advantage is robust to the specific proportion of causal loci that contribute genetic effects to a quantitative trait, which relates to trait architecture, as well as the evolutionary processes involved. We note that, as even more causal loci are added beyond the proportions explored in our study, the effects contributed by any individual locus becomes more diffuse, and global sample structure will become a more reasonable approximation of different causal loci with different local sample structures. In general, we found traits with “diffuse” genomic architecture (i.e., traits with a relatively high proportion of causal loci) to be challenging for all methods. Coal-Miner tended to cope better with the challenge relative to the other methods in our study, which we attribute to the design of the second stage in the Coal-Miner pipeline (i.e., candidate locus detection). Consistent performance trends were observed when comparing neutral versus non-neutral simulations. This suggests that, for the selection coefficients explored in our study, Coal-Miner’s performance is robust to the presence or absence of positive selection. A similar outcome was observed when comparing IUA model-based experiments involving two different types of model phylogenies – tree-like and non-tree-like.

The other model conditions in our simulation study explore alternative evolutionary scenarios where the proportion of causal loci was fixed. In these model conditions, Coal-Miner retained its performance advantage relative to the state-of-the-art, with one exception: Coal-Miner and Coal-Map had comparable AUROC on model conditions involving neutral evolution on non-tree-like model phylogenies and 10% causal loci, although Coal-Miner’s TPR at an FPR of 0.1 was a significantly better than Coal-Map’s. These model conditions involved the smallest proportion of causal loci in our study. We note that Coal-Map’s performance tended to degrade more rapidly than Coal-Miner as the proportion of causal loci increased, and the relative performance of the two methods may have changed for model conditions with higher proportions of causal loci that are otherwise equivalent.

Taken together, the model conditions included multiple sources of local genealogical variation, including genetic drift/ILS, gene flow, recombination, positive selection, and combinations thereof. We note that gene flow was not a necessary prerequisite for Coal-Miner’s performance advantage, so long as the other processes were involved (e.g., drift/ILS). The specific evolutionary processes contributing to local genealogical variation did not seem to matter as much as the presence of local genealogical variation, and Coal-Miner’s performance advantage was not necessarily predicated on specific evolutionary cause(s) of local genealogical discordance. These findings seem to suggest that Coal-Miner’s model and algorithm may be generalized to other evolutionary scenarios, so long as the breakpoint inference method used in stage one of the Coal-Miner pipeline suitable accounts for evolutionary processes with first-order contributions to genome evolution.

## 4.2 Empirical study

The empirical datasets in our study were more challenging than the simulated datasets because the former likely involved more complex evolutionary evolutionary scenarios compared to the latter. Additional evolutionary processes which may have played an important role include other types of natural selection and other demographic events (e.g., fluctuations in effective population size).

For both of the *Arabidopsis* datasets, Coal-Miner was able to detect significant associations in all five positive control regions. In contrast, neither GEMMA nor EMMAX – the statistical AM method used by Consortium [10] – were able to do the same. The vernalization requirement for flowering in *Arabidopsis* suggests that the flowering response at 16 °C presents a greater AM challenge than at 10 °C. Our findings were consistent with a need for more statistical power for the former as compared with the latter as well as the overall findings in the simulation study, which suggested that Coal-Miner offered improved statistical power relative to the state of the art. Coal-Miner also correctly analyzed positive and negative controls in the other empirical datasets. Furthermore, Coal-Miner analyses of the *Arabidopsis* and *Burkholdericeae* datasets identified putatively novel markers (i.e., markers which were not flagged using other AM methods). Additional comparative and functional analyses are needed to interpret these findings.

## 5 CONCLUSIONS

Across the range of genomic architectures and evolutionary scenarios explored in our study, Coal-Miner had comparable or typically improved statistical power and type I error control compared to state-of-the-art AM methods. The scenarios included different evolutionary processes such as genetic drift and ILS, positive selection, gene flow, and recombination – all of which can generate local genealogical variation that differs from the true species phylogeny. More work needs to be done to explore additional evolutionary processes which have first-order impacts on genome evolution (e.g., gene duplication and loss, other genome rearrangement events, etc.). As more divergent samples are included in a GWA study, more evolutionary processes potentially will become relevant to AM analysis. We fully expect that more algorithmic development will need to be done in this case, particularly regarding the breakpoint inference stage of Coal-Miner.

We conclude with our thoughts on future work. As an alternative to the pipeline-based design of Coal-Miner, simultaneous inference of local coalescent histories and AM model parameters will avoid error propagation across different stages of a pipeline-based algorithm. Furthermore, viewed through the lens of evolution, genotype and phenotype are arguably two sides of the same coin. The same could be said of “intermediate-scale” characters (e.g., interactomic characters). A combination of the extended coalescent models and LMMs could be used to capture evolutionary relatedness of and functional dependence between heterogeneous biological characters across multiple scales of complexity and at higher evolutionary divergences.

## Acknowledgements

The authors gratefully acknowledge the following support: National Science Foundation Grant CCF-1565719 (to KJL), a grant from the BEACON Center for the Study of Evolution in Action (NSF STC Cooperative Agreement DBI-093954) to KJL and GAB, and Michigan State University faculty startup funds (to KJL, to GAB, and to PPE). The authors would also like to thank the anonymous referees for their valuable feedback and suggestions.

